# H_2_O_2_ sulfenylates CHE linking local infection to establishment of systemic acquired resistance

**DOI:** 10.1101/2023.07.27.550865

**Authors:** Lijun Cao, Heejin Yoo, Tianyuan Chen, Musoki Mwimba, Xing Zhang, Xinnian Dong

## Abstract

In plants, a local infection can lead to systemic acquired resistance (SAR) through increased production of salicylic acid (SA). For 30 years, the identity of the mobile signal and its direct transduction mechanism for systemic SA synthesis in initiating SAR have been hotly debated. We found that, upon pathogen challenge, the cysteine residue of transcription factor CHE undergoes sulfenylation in systemic tissues, enhancing its binding to the promoter of SA-synthesis gene, *ICS1*, and increasing SA production. This occurs independently of previously reported pipecolic acid (Pip) signal. Instead, H_2_O_2_ produced by NADPH oxidase, RBOHD, is the mobile signal that sulfenylates CHE in a concentration-dependent manner. This modification serves as a molecular switch that activates CHE-mediated SA-increase and subsequent Pip-accumulation in systemic tissues to synergistically induce SAR.

**One Sentence Summary:** RBOHD-generated H_2_O_2_ sulfenylates transcription factor CHE to establish systemic acquired resistance in plants.

---

Systemic acquired resistance (SAR) is an inducible immune mechanism in plants (*1-3*). Unlike the animal adaptive immunity, SAR is triggered by local immune responses against specific pathogen effectors, known as effector-triggered immunity (ETI), and confers long-lasting protection to the rest of plants against a wide range of pathogens. This fascinating signaling phenomenon and its potential application in agriculture for engineering broad-spectrum disease resistance in crops have led to intense investigation and the identification of salicylic acid (SA) as the required signal for SAR (*4-7*). Though exogenous application of SA and its synthetic analogs has been shown to induce SAR without the ETI (*8-10*), for biological induction of SAR, how the local ETI induces systemic synthesis of SA has remained a mystery for more than 30 years. Multiple compounds, such as methyl salicylate (MeSA) (*11*), azelaic acid (AzA) (*12*), dehydroabietinal (DA) (*13*), glycerol-3-phosphate (G3P) (*14*) and pipecolic acid (Pip)/N-hydroxypipecolic acid (NHP) (*15-17*), have been identified as potential SAR signals that are involved in the production of SA and defense-related responses. However, a direct link between these mobile signals and systemic SA synthesis is missing.

## RESULTS

### CHE is a transcription factor required for SA synthesis and resistance only in systemic tissues

To identify the missing link between local ETI and systemic production of SA, we focused on the plant circadian clock transcription factor (TF), CCA1 HIKING EXPEDITION (CHE), which was found in our previous study to be required not only for the diurnal basal SA synthesis, but also for pathogen-induced SA production in systemic tissues (*18*). To better understand the regulation of CHE and how it contributes to the establishment of SAR over time, we performed time-course RNA-mediated oligonucleotide Annealing, Selection, and Ligation with next-generation sequencing (RASL-seq) (*19*) in both infected local and naïve systemic tissues. To minimize background variations between leaves, we used infected half-leaf as local tissues (Local) and the uninfected half-leaf as systemic tissues (Systemic), similar to the classic SAR experiment conducted by A. Frank Ross (*3*). After challenging *Arabidopsis thaliana* plants with the bacterial pathogen *Pseudomonas syringae* pv. *maculicola* (*Psm*) ES4326 carrying the effector gene *avrRpt2* (*Psm* ES4326/avrRpt2), the expression patterns of approximately 700 selected genes, with a particular focus on those involved in defense and SA production, were examined. We found similar overall gene expression patterns, as well as comparable levels of SA and Pip, in the *che* mutant compared to the wild type (WT) plants in the local tissues (Fig. 1, A and B, and fig. S1A). This finding is consistent with the previously reported normal local defense phenotype of the *che* mutant (*18*). However, in systemic tissues, the expression patterns were significantly different between WT and *che*, with the mutant showing lower gene inductions and substantial decreases in the production of SA and Pip (Fig. 1, A and B). These results show that TF CHE is required for SAR-associated gene expression and accumulation of SA and Pip only in systemic tissues. Interestingly, based on the RASL-seq data, the CHE-mediated systemic responses were not linked with significant transcriptional difference between WT and the *che* mutant of known TF genes *CALMODULIN BINDING PROTEIN 60G* (*CBP60g*) and *SAR DEFICIENT 1* (*SARD1*), which are involved in SA synthesis (*20*), *CIRCADIAN CLOCK-ASSOCIATED 1* (*CCA1*), which is a target clock gene of CHE (*21*), or *RESPIRATORY BURST OXIDASE HOMOLOGUE D* (*RBOHD*), which is known for immune-induced apoplastic ROS production (*22-26*) (fig. S1B).

**Fig. 1.**
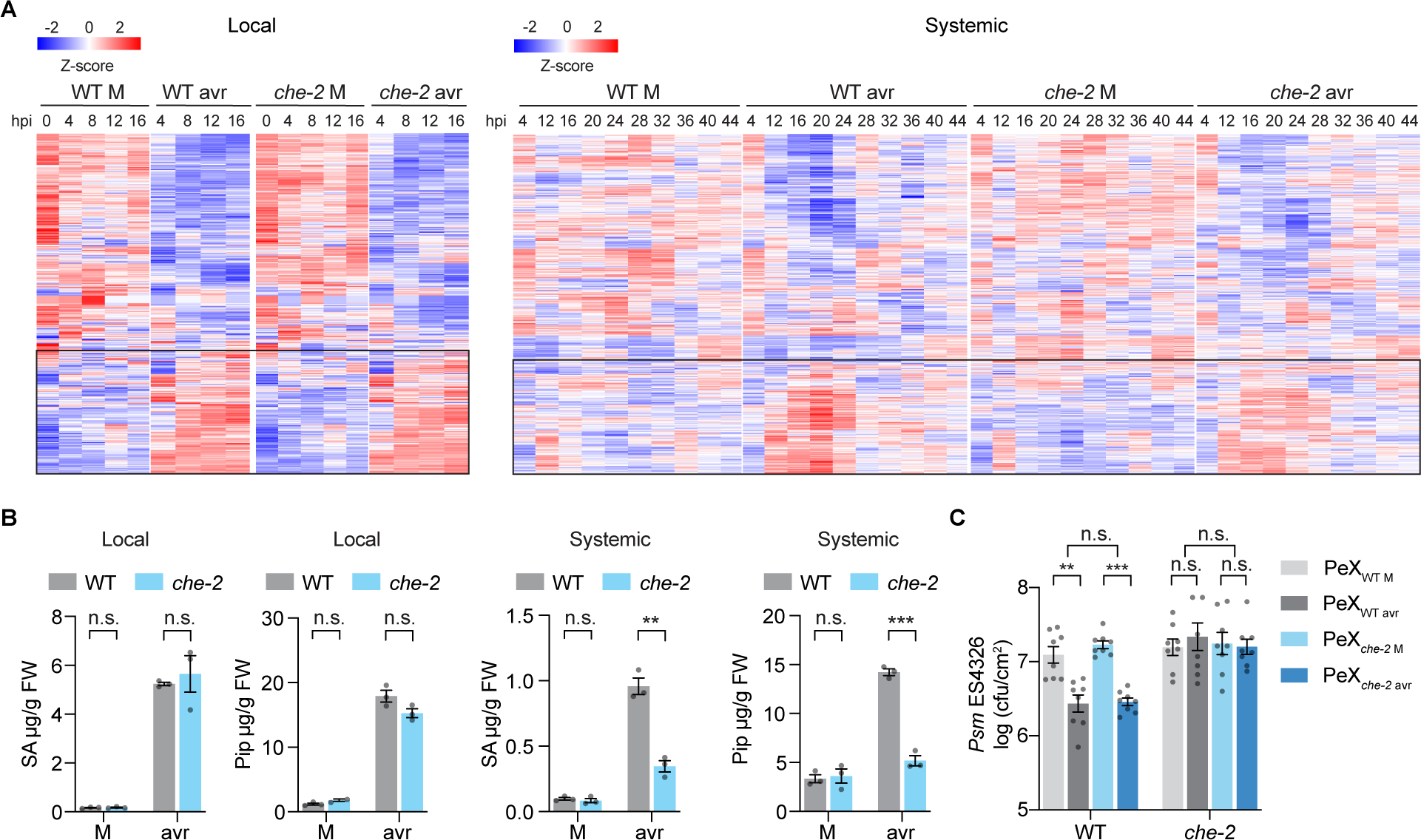
CHE functions specifically in the systemic tissues during SAR. (**A**) Heatmaps of normalized RASL-seq reads in local and systemic tissues after mock (M; 10 mM MgCl_2_) or *Psm* ES4326/avrRpt2 (avr; OD_600nm_ = 0.01) treatment. Black rectangles highlight the reads of upregulated genes by the avr treatment. hpi, hours post infiltration. (**B**) Levels of SA and Pip in local (12 hpi) and systemic tissues (24 hpi). Data are means ± SEMs (*n* = 3). (**C**) Bacterial growth in WT and *che-2* plants pre-treated with petiole exudate (PeX) collected from WT or *che-2* plants after M or avr treatment. Data are means ± SEMs (*n* = 8). Significant differences were calculated using either two-tailed Student’s t-tests or two-way ANOVA. \*\*\**P* < 0.001; \*\**P* < 0.01; \**P* < 0.05; n.s., not significant.

We next examined effects of the *che* mutant on the transportation and sensing of SAR-inducing signal(s) by measuring the SAR-inducing activity of petiole exudates (PeX) as previously described (*27-29*). Notably, PeX derived from WT and *che* mutant after *Psm* ES4326/avrRpt2 treatment exhibited similar capacities in reducing bacterial growth in WT plants (Fig. 1C). This observation suggests that the *che* mutant can produce SAR-inducing mobile signals like the WT. However, the inability of *che* mutant to restore defense through the application of PeX from either WT or *che* after pathogen challenge (Fig. 1C) indicates that CHE is defective in sensing the mobile signal. Consequently, these findings further confirm that CHE is specifically required for SA synthesis and resistance in systemic tissues.

### Cysteine mutants of CHE are defective in SAR

Since neither the *CHE* transcript nor the protein showed a significant increase in systemic tissues upon induction (fig. S2, A and B), we considered protein modification as an activation mechanism because CHE, also named TCP21, is one of the Class I TCP (TEOSINTE BRANCHED1, CYCLOIDEA, and PCF) TFs, which have a conserved cysteine residue in the non-canonical basic helix-loop-helix (bHLH) DNA-binding domain (*30*). Given that the cysteine-containing TCP proteins have been shown to be sensitive to redox conditions for their DNA-binding activity under different types of oxidants (*31*), and the cysteine residue of CHE is conserved across many plant species (fig. S2C), we decided to test whether mutating the single cysteine residue of CHE (cysteine 51) alters its function in SAR. We found that, unlike the WT *CHE* expressed by its native promoter (*CHEC*), the cysteine mutants *che^CS^* (cysteine to serine) and *che^CW^*(cysteine to tryptophan) failed in rescuing the SAR deficiency of the *che* mutants (Fig. 2, A and B, and fig. S3A) (*18*). This was accompanied by compromised systemic inductions of the SA synthesis gene *ISOCHORISMATE SYNTHASE 1* (*ICS1*), the SA-responsive gene *PATHOGENESIS-RELATED GENE 1* (*PR1*), and the Pip/NHP-synthesis genes *AGD2-LIKE DEFENSE RESPONSE PROTEIN 1* (*ALD1*) and *FLAVIN-DEPENDENT MONOOXYGENASE 1* (*FMO1*) (*32*) (fig. S3, B to E) and lower production of SA and Pip (Fig. 2, C and D) observed in the cysteine mutant line.

**Fig. 2.**
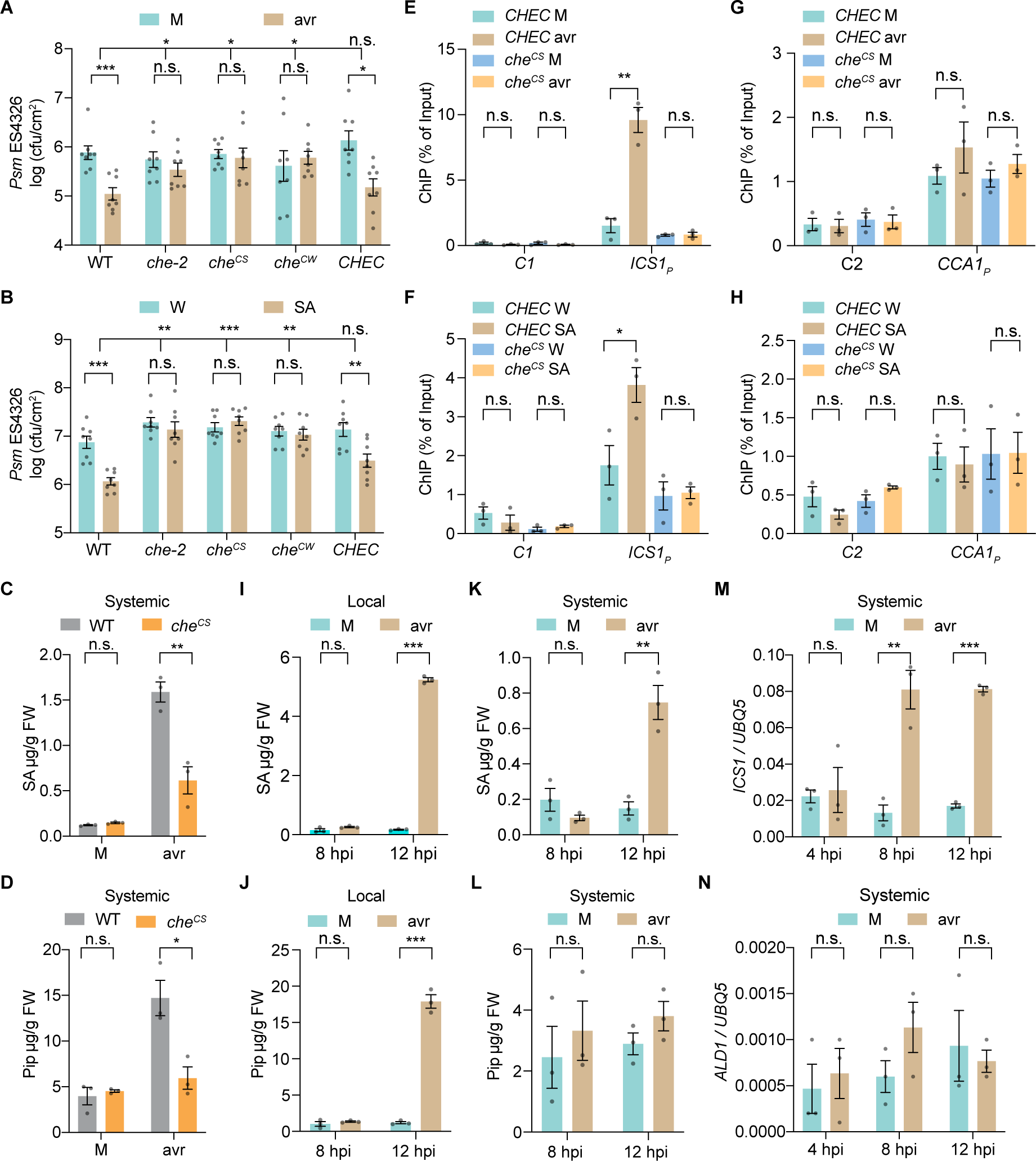
The cysteine residue in CHE is required for systemic SA synthesis, Pip accumulation, and SAR. (**A**) Bacterial growth in systemic tissue after local pathogen induction. Plants were infiltrated with mock (M; 10 mM MgCl_2_) or *Psm* ES4326/avrRpt2 (avr; OD_600nm_ = 0.01) 2 days before infiltration of the systemic tissues with *Psm* ES4326 (OD_600nm_ = 0.001), and bacterial growth was measured 3 days after the second infiltration. Data are means ± SEMs (*n* = 8). *CHEC*, *che-2* transformed with the WT *CHE* expressed by its native promoter. *che^CS^* and *che^CW^*, transformants expressing the cysteine-to-serine and cysteine-to-tryptophan mutant *che*, respectively. (**B**) Bacterial growth after SA treatment. Plants were sprayed with water (W) or 1 mM SA 1 day before infiltration of *Psm* ES4326, and bacterial growth was measured 3 days after the infiltration. Data are means ± SEMs (*n* = 8). (**C** and **D**) Levels of SA (C) and Pip (D) in systemic tissues after local M or avr treatments. Data are means ± SEMs (*n* = 3). (**E** and **F**) ChIP-qPCR analysis of CHE binding to the *ICS1* promoter carrying the TCP-binding site (*ICS1_P_*) using systemic tissues after local M or avr treatment (E) and W- or SA-treated tissues (F). *C1*, control for *ICS1p*. Data are means ± SEMs (*n* = 3). (**G** and **H**) ChIP-qPCR analysis of CHE binding to the *CCA1* promoter carrying the TCP-binding site (*CCA1_P_*) using systemic tissues after local M or avr treatment (G) and W- or SA-treated tissues (H). *C2*, control for *CCA1p*. Data are means ± SEMs (*n* = 3). (**I** and **J**) Time-course measurement of SA (I) and Pip (J) in the local tissues. Data are means ± SEMs (*n* = 3). hpi, hours post infiltration. (**K** and **L**) Time-course measurement of SA (K) and Pip (L) in the systemic tissues. Data are means ± SEMs (*n* = 3). (**M** and **N**) Time-course transcriptional level of *ICS1* (M) and *ALD1* (N) in the systemic tissues of WT plants. Data are means ± SEMs (*n* = 3). Significant differences were calculated using either two-tailed Student’s t- tests or two-way ANOVA. \*\*\**P* < 0.001; \*\**P* < 0.01; \**P* < 0.05; n.s., not significant.

To examine the effect of the cysteine-mutant on the transcriptional activity regulated by CHE, we performed chromatin immunoprecipitation-quantitative PCR (ChIP-qPCR) analysis and found that after pathogen challenge, the binding of CHE to the *ICS1* promoter (*ICS1_p_*), which carries the TCP-binding site, was significantly increased in the systemic tissues of *CHEC* transgenic plants, while this increase was compromised in the transgenic plants expressing che^CS^ (Fig. 2E). We also performed exogenous SA treatment and observed a similar defect in SA-induced promoter-binding for the che^CS^ mutant protein (Fig. 2F), consistent with our previous work showing that the *che* mutants are defective in SA-mediated signal amplification (*18*). However, the binding of CHE to the clock gene *CCA1* promoter (*CCA1p*), which also carries a TCP-binding site (*21*), was not significantly increased after pathogen challenge (Fig. 2G) nor after SA treatment (Fig. 2H). Consistent with our observations that the che^CS^ mutant could rescue the short period of *CCA1* expression (*21*) in the *che-1 lhy-20* double mutant (fig. S4, A and B). These data indicate that CHE regulates systemic SA synthesis independent of its clock function. Furthermore, we found that CHE does not bind to the promoter of *ALD1* or *FMO1* (fig. S5, A and B), suggesting that the regulation of SAR by CHE is not directly through these genes. Taken together, these results demonstrate that the cysteine residue of CHE regulates SA synthesis in systemic tissues through the modulation of its binding to the *ICS1* promoter.

### CHE-mediated SA synthesis in systemic tissues occurs prior to and independent of the Pip increase

SA and Pip are both important molecules that regulate SAR upon pathogen infection (*4, 16, 32*) and the systemic induction of both molecules are compromised in *che* mutants (Fig. 1B, and Fig. 2, C and D). Pip is one of the early accumulated signals and it was shown to function upstream of many other SAR-inducing signals, including AzA and G3P, upon pathogen challenge (*33-35*). Therefore, it is important to know whether Pip is induced earlier than SA during the initiation of SAR. We performed a time-course measurement of SA and Pip levels in both local and systemic tissues. The results showed that SA and Pip were significantly increased in the local tissues 12 hours post pathogen infiltration (Fig. 2, I and J). However, in the systemic tissues, at 12 hours post pathogen infiltration, SA level was elevated (Fig. 2K), whereas Pip was not (Fig. 2L), suggesting that the initial systemic induction of SA might not be through the Pip pathway, consistent with a previous report that *de novo* synthesis of Pip in systemic tissues is dependent on SA (*33*). Moreover, induction of the SA synthesis gene *ICS1* was observed in systemic tissues at 8 hours post pathogen infiltration, whereas an increase in the Pip synthesis gene *ALD1* was not detected even at 12 hours post pathogen infiltration (Fig. 2, M and N). These results indicate that systemic inductions of *ICS1* and SA production by CHE occur prior to Pip induction. In our analysis, the Pip derivative NHP was not detected despite that we successfully detected NHP standard, probably due to instability of NHP as suggested by previous studies (*15, 36*).

### CHE is sulfenylated in systemic tissues after a local induction of ETI

To investigate how CHE initiates SAR, we focused on its functionally important cysteine residue. The thiol moiety (-SH) of cysteine residue is highly reactive and can undergo various modifications by reactive oxygen species (ROS), including forming disulfide bond between two cysteines, S-sulfenylation (SOH), S-sulfinylation (SO_2_H) and S-sulfonylation (SO_3_H) (*37*). Since CHE has only one cysteine residue, a disulfide bond formation would require a cysteine-containing partner which was not detected under either mock or induced conditions using non-reducing SDS gel electrophoresis (fig. S2B). We then performed a biotin-switch assay (*38*) to examine other possible modifications. Excitingly, we found that CHE is specifically sulfenylated (CHE-SOH) in the systemic tissues between 8 and 36 hours post infiltration with *Psm* ES4326/avrRpt2, but not from mock-treatment (Fig. 3A, and fig. S6A). In addition, we found CHE in the distal systemic tissues also underwent sulfenylation when lower leaves were challenged with *Psm* ES4326/avrRpt2 (fig. S6B). This confirms that both the adjacent and distal systemic tissues employ similar mechanism in establishing SAR as suggested in the classic study (*3*). In addition, exogenous application of SA could also lead to CHE-SOH, though later at 24 hours after the treatment (Fig. 3A). To determine whether CHE can be sulfenylated *in vitro*, we treated purified CHE with various concentrations of H_2_O_2_, as H_2_O_2_ is known to be produced during plant defense responses including ETI (*39, 40*). Interestingly, we found that CHE is sulfenylated specifically at 50 μM H_2_O_2_ treatment, while lower or higher concentrations did not result in sulfenylation (Fig. 3B).

**Fig. 3.**
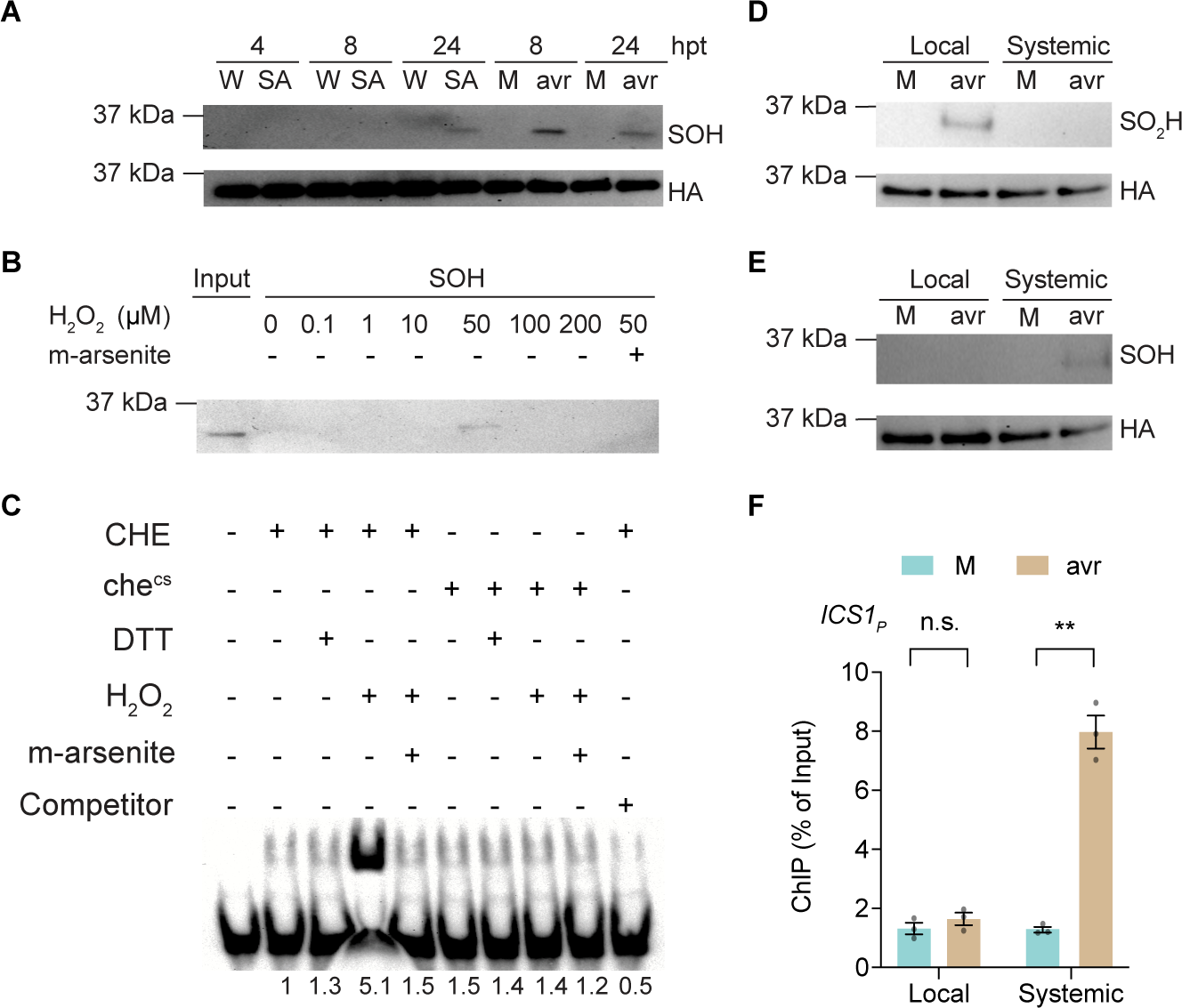
Sulfenylation of CHE occurs specifically in systemic tissues during SAR. (**A**) Sulfenylation (SOH) of CHE in systemic tissues after local mock (M; 10 mM MgCl_2_) or *Psm* ES4326/avrRpt2 (avr; OD_600nm_ = 0.01) treatment, or in the tissues after water (W) or 1 mM SA treatment. hpt, hours post treatment. (**B**) *In vitro* sulfenylation of CHE after H_2_O_2_ treatment and specific removal of sulfenylation by 50 mM sodium arsenite (m-arsenite). (**C**) Electrophoresis mobility shift assay of recombinant CHE protein binding to the DNA probe of TCP-binding site of *ICS1* promoter (*TBSi*). (**D** and **E**) Sulfinylation (SO_2_H) (D) and SOH (E) of CHE in local and systemic tissues, respectively, after M or avr treatment. Samples are collected 7 to 8 hours post pathogen treatment. (**F**) ChIP-qPCR analysis of CHE binding to the *ICS1* promoter carrying the TCP-binding site (*ICS1_P_*) in the local and systemic tissues after M or avr treatments. Data are the means ± SEMs (*n* = 3). Significant differences were calculated using two-tailed Student’s t-tests. \*\**P* < 0.01; n.s., not significant.

To determine whether the *in vivo* CHE-binding to the *ICS1* promotor (Fig. 2E) is due to sulfenylation of its cysteine residue, we performed a gel electrophoresis mobility shift assay using a DNA probe containing the TCP-binding site from the *ICS1* promoter (*TBSi*). We found that treatment with 50 μM H_2_O_2_ significantly enhanced the binding of CHE to *TBSi in vitro* (Fig. 3C). This increase in promoter-binding is specific to CHE-SOH because treatment of H_2_O_2_-induced CHE with the specific reducing agent for the sulfenic acid, sodium arsenite (m-arsenite), diminished the binding, similar to the general reducing agent DTT treatment (Fig. 3, B and C). Moreover, the H_2_O_2_-induced binding was largely absent in the che^CS^ mutant protein (Fig. 3C), which further confirmed that sulfenylation of the cysteine residue in CHE is the molecular switch that increases its binding to the *ICS1* promoter.

To address the question why CHE sulfenylation is only detected in systemic tissues, we hypothesized that, as suggested by our H_2_O_2_-concentration-dependent sulfenylation experiment (Fig. 3B), the higher ROS levels in the local tissues might cause further oxidation of the cysteine residue in CHE. We found that this was indeed the case because sulfinylation (CHE-SO_2_H) was detected in the local tissues while sulfenylation (CHE-SOH) in systemic tissues (Fig. 3, D and E). Regardless of whether CHE-SO_2_H in the local tissues can be further oxidized to CHE-SO_3_H, neither form is active in binding to the *ICS1* promoter as shown by the ChIP-qPCR results (Fig. 3F). This H_2_O_2_-concentration-specific activation of CHE also explains the different conclusions on the role of H_2_O_2_ observed in previous studies which utilized different H_2_O_2_ concentrations (*41, 42*).

### Production of H_2_O_2_ by RBOHD during ETI serves as a mobile signal for SAR

Since *in vitro* treatment of CHE protein by H_2_O_2_ can cause its sulfenylation, we investigated the possibility of H_2_O_2_ serving as the endogenous signal for activating CHE by measuring H_2_O_2_ levels in both local and systemic tissues after a local ETI induction. As expected, we detected higher induction of H_2_O_2_ levels in the local tissues than in the systemic tissues from 4 hours post pathogen infiltration (Fig. 4A, and fig S7, A and B). The increased production of H_2_O_2_ in distal systemic tissues starts around 8 hours post pathogen challenge in the lower leaves (fig. S7C). Interestingly, this H_2_O_2_ increase was not significantly affected in the *che* mutant, except at 24 hours post pathogen infiltration in the systemic tissues (Fig. 4A, and fig S7, A and B), nor in the *ald1* or *fmo1* mutants; but abolished in *rps2,* the receptor *RESISTANT TO P. SYRINGAE 2* mutant for the bacterial effector avrRpt2 (*43, 44*); in *sid2,* a *SALICYLIC ACID INDUCTION DEFICIENT 2* mutant in the SA-synthesis gene *ICS1 (45*); and in *rbohD*. Consistently, H_2_O_2_ levels in petiole exudates collected from the *che*, *ald1* and *fmo1* mutant plants were similarly increased as in the WT, while those from the *rbohD* and *rps2* mutants showed no increase (Fig. 4B). These results demonstrate that the production of H_2_O_2_ during defense does not require the previously reported mobile signals Pip or NHP (*15, 17, 33*) but requires RBOHD and the receptor for the effector RPS2. Moreover, SA treatment could also induce production of H_2_O_2_ to a level similar to that detected in systemic tissues induced by ETI (fig. S7D). This indicates the involvement of SA in the production of H_2_O_2_, which is consistent with the previous reports (*34, 41*) and the observation that ETI-induced H_2_O_2_ production is compromised in the SA-deficient *sid2* mutant (fig. S7, A and B).

**Fig. 4.**
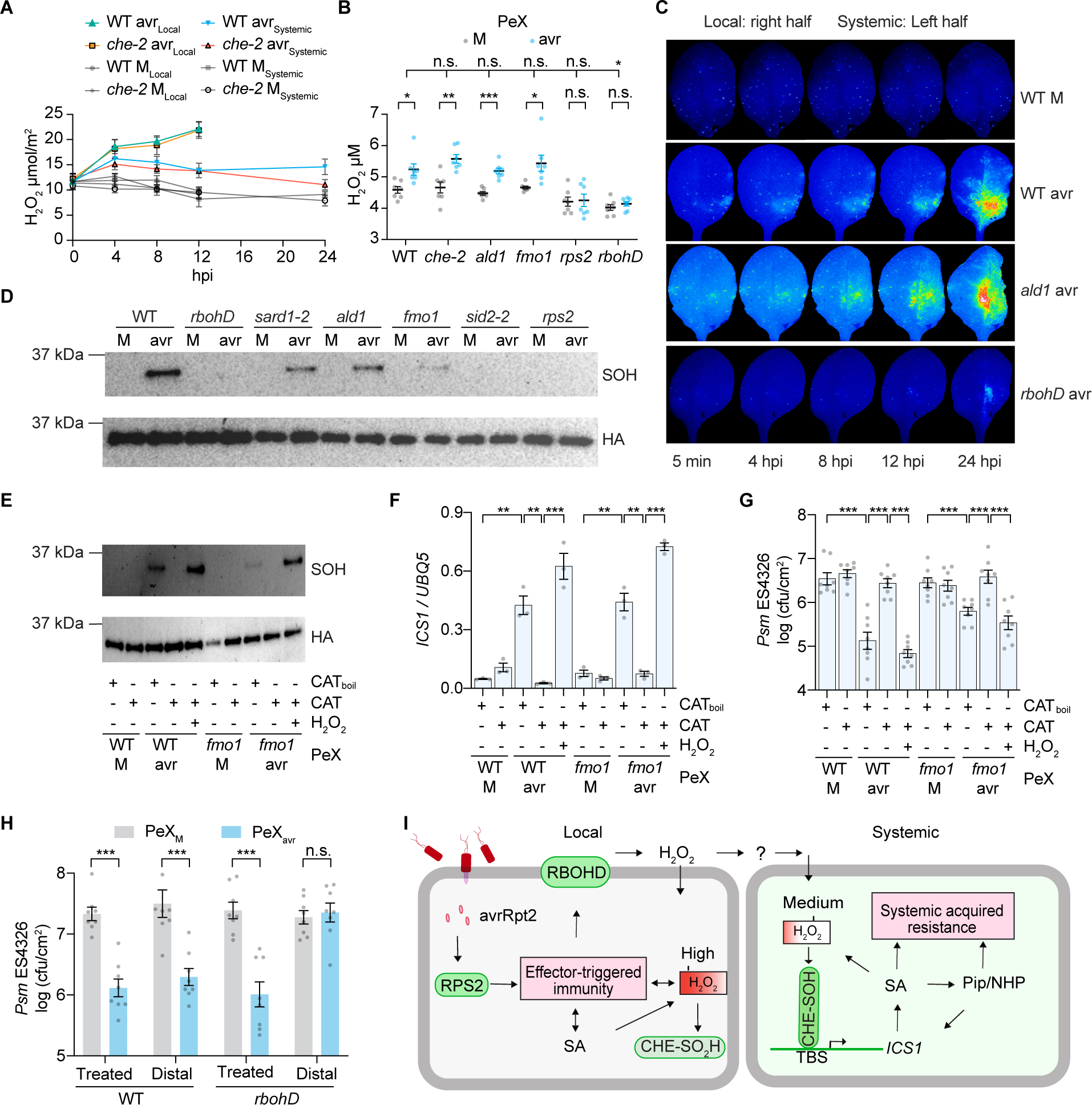
H_2_O_2_ produced by RBOHD is the mobile signal that sulfenylates CHE to activate systemic SA synthesis and resistance. (**A**) Time-course measurement of H_2_O_2_ in local and systemic tissues of plants after mock (M; 10 mM MgCl_2_) or *Psm* ES4326/avrRpt2 (avr; OD_600nm_ = 0.01) treatment. Data are means ± SEMs (*n* = 5). hpi, hours post infiltration. (**B**) H_2_O_2_ levels in petiole exudates (PeX) collected after M or avr treatment. Data are means ± SEMs (*n* = 7). (**C**) Snap shots from live imaging of ROS production and transportation using 2’,7’- dichlorodihydrofluorescein diacetate after treatment with M or avr (also see Movies S1-3). (**D**) Sulfenylation (SOH) of CHE in systemic tissues after local M or avr treatment in different genetic backgrounds. (**E**-**G**) The effect of catalase treatment of petiole exudates collected after M or avr treatment on CHE sulfenylation (E), *ICS1* expression (F), and bacterial growth (G). Data are means ± SEMs, *n* = 3 for (F), *n* = 8 for (G). (**H**) Bacterial growth in the treated or distal leaves of WT or *rbohD* plants after treatment with PeX collected from WT plants after M or avr treatment. Data are means ± SEMs (*n* = 8). (**I**) The working model of CHE-mediated SAR induction. The perception of bacterial effector avrRpt2 by the immune receptor RPS2 induces ETI and local production of SA, resulting in accumulation of high H_2_O_2_ concentration and CHE sulfinylation (CHE-SO_2_H), which lacks the increased binding activity to the *ICS1* promoter. Local apoplastic H_2_O_2_ produced and transported/relayed via an unknown mechanism (?) by RBOHD to systemic tissues to sulfenylate CHE (CHE-SOH). CHE-SOH binds to *ICS1* promoter to induce *de novo* SA synthesis and initiate SAR. Full-scale resistance is achieved through an amplification loop involving synergistic activities of SA and Pip/NHP. Significant differences were calculated using either two-tailed Student’s t-tests or two-way ANOVA. \*\*\**P* < 0.001; \*\**P* < 0.01; \**P* < 0.05; n.s., not significant.

The requirement of the RPS2 immune receptor for the production of H_2_O_2_ during SAR is noteworthy because while induction of SAR was initially reported to be associated with necrosis in the local tissues (*3, 46*), there have been reports showing induction of systemic resistance by virulent pathogens that do not trigger visually detectable cell death (*16, 47-49*). Therefore, we used the type III secretion system mutant *Ps pv. tomato* (*Pst*) DC3000 *hrcC^-^*, which is unable to deliver effectors to the plant host (*50*), to examine the sulfenylation of CHE in systemic tissues. Our results revealed that, under the same inoculation conditions, *Pst* DC3000 *hrcC^-^* failed to induce systemic CHE-SOH or the expression of *ICS1* or *PR1,* unlike the *Pst* DC3000/avrRpt2 control (fig. S7, E to G). These findings suggest that a strong and sustained production of H_2_O_2_ in local tissues is required for the establishment of SAR.

To monitor the transportation of ROS, including H_2_O_2_, from the local to systemic tissues during SAR induction, we employed a live imaging technique using 2’,7’- dichlorodihydrofluorescein diacetate (*51, 52*). Upon pathogen challenge, strong ROS signals were detected in both WT and *ald1* plants in the local tissues, and the transportation of ROS signal(s) to systemic tissues was subsequently observed in both genetic backgrounds (Fig. 4C, and Movies S1 and S2), indicating that ROS (e.g., H_2_O_2_) signaling is not directly dependent of production of Pip/NHP. Moreover, the induced ROS production occurred earlier following pathogen challenge in both local and systemic tissues than other reported mobile signals, such as G3P, AzA, DA and Pip (*32, 35*) (Fig. 4C, and Movies S1 and S2). However, we observed a marked reduction in ROS production and transportation in the *rbohD* mutant (Fig. 4C, and Movie S3), consistent with the diminished H_2_O_2_ levels observed in this mutant in our quantitative measurements (Fig. 4B, and fig. S7, A and B). RBOHD is known to catalyze the production of H_2_O_2_ by transfer electrons to oxygen to form superoxide anion (O_2-_) which is subsequently converted to H_2_O_2_ through enzymatic or non-enzymatic dismutation (*53*). Furthermore, in the *rbohD* mutant, we observed a deficiency of CHE-SOH (Fig. 4D) and reduced CHE binding to the *ICS1* promoter (fig. S8A), as well as compromised induction of *ICS1* and systemic-induced resistance to *Psm* ES4326 (fig. S8, B and C). Based on these findings, we hypothesized that RBOHD-produced H_2_O_2_ during ETI is a mobile signal for the establishment of SAR.

### H_2_O_2_ in petiole exudates is responsible for sulfenylation of CHE, SA synthesis, and induction of resistance in systemic tissues

We next examined whether H_2_O_2_ is the signal that sulfenylates CHE and establishes SAR in naïve tissues. We found that pretreating the petiole exudates collected from WT and the *fmo1* mutant with the H_2_O_2_-scavenging enzyme, catalase (CAT), led to depletion of H_2_O_2_ (fig. S8D). This treatment also inhibited the exudate’s ability in sulfenylating CHE (Fig. 4E) and inducing the expression of *ICS1* (Fig. 4F), *ALD1*, *FMO1* and *PR1* (fig. S8, E to G), and in triggering resistance against *Psm* ES4326 (Fig. 4G). This inhibitory effect of catalase was abolished by heat denaturation (CAT_boil_) and could be overcome by exogenous application of H_2_O_2_ after removing the enzyme (Fig. 4, E to G, and fig S8, E to G). These results further support our hypothesis that H_2_O_2_ production mediated by RBOHD is the initiating mobile signal that activates CHE through sulfenylation to induce SAR. It is interesting that, compared to the petiole exudates collected from WT, those collected from the *fmo1* mutant were less active in inducing *ALD1* and *FMO1* genes (fig. S8, E and F) and in conferring resistance to *Psm* ES4326 (Fig. 4G), consistent with the reduced sulfenylation of CHE in *sard1,* a mutant that affects SA and Pip synthesis (*20, 54*), in *ald1* and *fmo1* mutants (Fig. 4D). These results suggest the existence of mutual amplifications between SA and Pip/NHP signaling in systemic tissues which is consistent with the previous reports that Pip/NHP, together with SA, play an important role in the establishment of SAR (*32*). It is interesting that while PeX collected from WT after pathogen challenge-initiated defense against *Psm* ES4326 in both the treated and distal leaves of WT plants, it only triggered defense in the treated leaves of *rbohD* plants (Fig. 4H), indicating that distal SAR requires the transport or relay-production of H_2_O_2_ by this membrane-associated NADPH oxidase.

In summary, H_2_O_2_ produced through RBOHD during ETI is the primary signal that sulfenyaltes CHE and induces the expression of *ICS1*, which directly results in *de novo* synthesis of SA in systemic tissues. H_2_O_2_-mediated sulfenyaltion of CHE is required for SA and Pip accumulation in the systemic tissues and both signals function synergistically to fully induce SAR (Fig. 4I).

## Discussion

It is intriguing that a circadian clock TF (i.e., CHE) is used to control SAR in plants. It raises questions about the biological significance of this phenomenon. Since the redox-sensitive cysteine residue in CHE is highly conserved among plant species (fig. S2C), it is likely that H_2_O_2_-mediated sulfenylation and activation of CHE for SAR is a conserved mechanism in plants. Interestingly, the mutation of the cysteine residue in CHE does not affect its clock function (Fig. 2, G and H, and fig. S1B and S4), indicating that the crucial role of CHE in SAR is a novel function beyond circadian clock regulation. Therefore, the significance of CHE’s role in SAR probably lies upon its own diurnal expression, which peaks around dusk (*18, 55*) (fig. S2A). Indeed, we have shown in a separate study that ETI-associated cell death is gated by the redox rhythm towards the morning (*56*). Therefore, the peak expression of CHE around dusk coincides with its sulfenylation at 8-12 hours after the morning induction of ETI (Fig. 3A) and also consistent with CHE-mediated basal SA synthesis which occurs at night and reaches the maximum before dawn (*18*).

The discovery of H_2_O_2_ produced during ETI as the initiating mobile signal and H_2_O_2_- mediated sulfenylation of CHE as the signal transduction mechanism explains “the classic” SAR (*1-3, 57*) because of the requirement of RPS2-mediated ETI for the sulfenylation of CHE in systemic tissues (Fig. 4D, fig. S7, E to G). However, studies have shown that infection by virulent pathogens can also induce enhanced disease resistance in plant systemic tissues (*2, 48, 49*). Whether these responses involve distinct induction mechanisms require further investigation. It is conceivable that a long-lasting H_2_O_2_ production during a local induction event is the main determining factor. Whereas microbe-associated molecular patterns (MAMPs)-treatment can activate RBOHD and trigger apoplastic H_2_O_2_ production (*25, 53*), pathogen-induced ETI, which involves the detection of both MAMP and effector signals, can lead to a strong and prolonged H_2_O_2_ production (*58*). The more transient nature of the H_2_O_2_ increase during PTI might explain the lack of systemic CHE sulfenylation and defense gene expression after a local inoculation by *Pst* DC3000 *hrcC^-^*, in contrast to the *Pst* DC3000/avrRpt2 (fig. S7, E to G). Therefore, the dose of pathogen inoculant and the testing time after the initial induction may all have an impact on the result. Our finding that an optimal concentration of H_2_O_2_ is required to sulfenylate CHE might also reconcile conflicting reports on the role of H_2_O_2_ in defense (*41, 42*) because if a higher-than-optimal level of H_2_O_2_ were used in treating plants, CHE would be inactivated through further oxidation of the cysteine (Fig. 3, B to F, and 4A). It is tempting to hypothesize that the sensitivity of CHE to H_2_O_2_ concentration would allow plants to gauge the level of local infection to determine whether and how much to activate SAR in naïve tissues. Therefore, our study not only identifies H_2_O_2_ as the initiating mobile signal and H_2_O_2_-concentration-dependent sulfenylation of CHE in inducing *de novo* systemic SA synthesis as the signal transduction mechanism, but also conforms with many discoveries made in the past 30 more years to show how a local infection modulates the level of SAR (*1-3, 32*).

## Acknowledgments

We thank Dr. Sheng Yang He for *Pst* DC3000 *hrcC^-^* and *Pst* DC3000/avrRpt2 bacteria strains; Dr. Jose Pruneda-Paz for WT (*CCA1_P_:LUC*), *che-1* (*CCA1_P_:LUC*), *lhy-20* (*CCA1_P_:LUC*) and *che-1 lhy-20* (*CCA1_P_:LUC*) mutant lines. We thank Dr. Pei Zhou, Mingru Li and Dr. Christoph F. Schmidt for help in measuring H_2_O_2_ levels. We also thank Lucas Li at the Duke Proteomics and Metabolomics Core Facility for measuring SA and Pip.

## Funding

National Institutes of Health grant NIH 1R35GM118036 (XD)

National Science Foundation grant NSF IOS-1645589 and IOS-2041378 (XD)

Howard Hughes Medical Institute (XD)

Hargitt Postdoctoral Fellowship (LC, HY)

## Author contributions

Conceptualization: LC, HY, XD

Methodology: LC, TC, XD

Investigation: LC, HY, TC, MM, XZ

Funding acquisition: LC, HY, XD

Supervision: XD

Writing – original draft: LC, XD

Writing – review & editing: LC, HY, TC, XZ, XD

## Competing interests

XD is a founder of Upstream Biotechnology Inc. and a member of its scientific advisory board, as well as a scientific advisory board member of Inari Agriculture Inc. and Aferna Bio.

## Data and materials availability

All generated materials in the manuscript are available upon request. All data are available in the main text or the supplementary materials.

## Supplementary Materials

Materials and Methods

Figs. S1 to S8

Tables S1 to S3

Movies S1 to S3

## Materials and Methods

### Plant materials

*Arabidopsis thaliana*, including wild type (WT), *che-2* (*21*), *che-1* (*21*), *che-3* (SALK_106694), *lhy-20* (*21*), *ald1 (59)*, *fmo1 (60)*, *rps2* (*43, 44*), *sid2-2 (45)*, *rbohD (61)*, *sard1-2* (*20*), and other transgenic plants used in this study are all in the Columbia-0 ecotype background. Plants were grown in soil at 22 °C under 12 hours light/12 hours dark with 60% relative humidity.

### Plasmid construction and plant transformation

To generate CHE complementation lines (*CHEC*), the *CHE* native promoter driving the coding sequence was used to transform *che-2*. Fragments were PCR amplified from the genomic DNA and cloned into pDONR207 with Gateway^TM^ BP Clonase^TM^ II Enzyme Mix (ThermoFisher Scientific) and then cloned into pGWB513 with Gateway^TM^ LR Clonase^TM^ II Enzyme Mix (ThermoFisher Scientific) for plant transformation. Vectors of cysteine mutants (cysteine mutated to serine in *che^CS^* and cysteine mutated to tryptophan in *che^CW^*) were generated through PCR amplification with corresponding primers (Table S1) using pGWB513 carrying *CHEC* as the template, then the *CHEC* template vector were removed by digestion with DpnI (NEB) and the cysteine mutant vectors were recovered by re-transformation in DH5α competent cells. WT or *che^CS^* coding sequence was constructed into pGEX6P1-DEST-HA and used for protein expression and purification in *E. coli*. Native promoter-driven the coding sequence of WT or *che^CS^* cloned into pGWB513 was used for transformation of *che-1* (*CCA1_P_:LUC*) and *che-1 lhy-20* (*CCA1_P_:LUC*) mutants. Native promoter-driven WT CHE coding sequence cloned into pGWB513 was used for transformation of *rbohD*, *sard1-2*, *ald1*, *fmo1*, *sid2-2*, and *rps2* mutants. All plasmids constructed were confirmed by Sanger sequencing before use. *Agrobacterium*-mediated transformation was performed as described (*62*).

### RASL-seq

Total RNA extracted using TRIzol (Ambion) was processed as described (*19*). Primers used in RASL-seq are listed in Tables S2 and S3. The raw sequencing data were first aligned to gene-specific primers. The gene counts were subjected to library size normalization and log2 transformation. The resulting gene expression matrix is provided in Tables S2 and S3.

### Quantitative real-time PCR

Total RNA was extract from leaf tissues of 3-week-old plants using TRIzol, followed by DNase I (Invitrogen) treatment to remove genomic DNA contamination. The extracted RNA was then used as a template for reverse transcription using SuperScript^®^ III Reverse Transcriptase (Invitrogen) with an oligo (dT) primer. FastStart Universal SYBR Green Master (Roche) was used in real-time PCR with primers listed in Table S1. *Ubiquitin 5* (*UBQ5*) was used as the internal control.

### SA and Pip quantification

SA and Pip were measured as described (*17*) with modifications. 3-6 leaves were collected, weighed, and ground in liquid nitrogen. The samples were then extracted in 1 mL of MeOH/H_2_O (80/20, v/v), then vortexed for 1 min and incubated at 4 °C for 10 min with rotation. The supernatant was collected after centrifugation at 4 °C (14,000 g for 10 min). The extraction was repeated one more time before combining the two-extracted supernatants into a 2-mL tube, then the supernatant was collected after another centrifugation. 400-800 μL of the extract was evaporated to dryness using an Eppendorf concentrator plus/Vacufuge^®^ plus system. Then, 100 μL of a 50 % methanol solution was added into the sample vials and vigorously vortexed. The solution was subjected to centrifugation at 5 °C (15,000 rpm for 5 min) and 40 μL of the resulting supernatant was transferred into the well of a 1 mL 96-well NUNC plate (ThermoFisher Scientific). After adding 10 μL internal mixture (0.5 μg/mL salicylic acid-d_4_ and 0.5 μg/mL pipecolic acid-d_9_), the plate was mixed on a ThermoMixer at 1,000 rpm for 10 min, then centrifuged at 3,000 rpm for 2 min before injection into the instrument.

Samples were analyzed with the Waters TQ-X MS system (Milford, MA) with the Acquity UPLC. Software Masslynx 4.2 was used for data acquisition. The LC separation was performed on a Waters Acquity CSH phenyl-hexyl column (2.1 x 100 mm, 1.7 μm) with mobile phase A (0.1 % formic acid in water) and mobile phase B (0.1 % formic acid in acetonitrile). The flow rate was 0.45 mL/min. The linear gradient was as follows: 0-1 min, 100 % A; 3 min, 50 % A; 3.1-4.5 min, 0 % A; 4.6-6.1 min, 100 % A. The autosampler was set at 10 °C and the column was kept at 45 °C. The injection volume was 2 μL. Mass spectra were acquired under both positive and negative electrospray ionization (ESI) with MRM as the detection approach. The positive mode was used for pipecolic acid (m/z 130.0 → m/z 67.0) and internal standard pipecolic acid-d_9_ (m/z 139.0 → m/z 61.0). The negative mode was used for salicylic acid (m/z 137.0 → m/z 93.0) and internal standard salicylic acid-d_4_ (m/z 141.0 → m/z 97.0).

### Petiole exudate collection and treatment

The petiole exudate (PeX) was collected as previously described (*63*) with modifications. Specifically, we first tried two different concentrations of *Psm* ES4326/avrRpt2 (OD_600nm_ = 0.01 and OD_600nm_ = 0.0005) to inoculate 3-week-old plant leaves, and excised the leaves at 8 and 24 hours, respectively, for PeX collection for 48 hours. We found that after proper dilution, both inoculants produced similar results in H_2_O_2_ production, CHE sulfenylation and plant defense against *Psm* ES4326. Consequently, we used OD_600nm_ = 0.0005 to simulate a natural infection more closely for the experiments. Another change made in this study was the inclusion of 0.02 % acetanilide in the collection solution to stabilize H_2_O_2_. The collected PeXs were then used to treat plants, from which leaf samples were collected 4 hours after PeX treatment for protein sulfenylation analysis or 24 hours after PeX treatment for bacterial growth and quantitative real-time PCR analysis. 50 μl of the PeX was used to quantify H_2_O_2_ concentration. To remove H_2_O_2_ from the PeX, 0.1 mg/mL catalase (Sigma) was used. To denature catalase, the catalase solution was incubated at 95 °C for 15 min. To rescue the catalase-treated PeX, an Amicon^®^ Ultra-4 centrifugal filter (Millipore) was used to remove the enzyme, and 10 μM H_2_O_2_ was added to the collected PeX.

### ETI-induction of SAR and measurement of bacterial growth

SAR was induced by infiltrating half-leaf with *Psm* ES4326/avrRpt2 (OD_600nm_ = 0.01) as the local tissues and the other un-infiltrated half-leaf was used as the systemic tissues (except specific mentioned) for RASL-seq, SA and Pip production quantifications, quantitative real-time PCR, the sulfenylation and sulfinylation of CHE, chromatin immunoprecipitation, H_2_O_2_ production assay, and ROS production assays. Plants were also infiltrated with *Pst* DC3000 *hrcC^-^* (OD_600nm_ = 0.01) and *Pst* DC3000/avrRpt2 (OD_600nm_ = 0.01) using half-leaf and samples were collected 24 hours after infiltration for the sulfenylation of CHE and quantitative real-time PCR assays.

For examining SAR protection against bacterial growth, 3-week-old plants were first infiltrated with 10 mM MgCl_2_ (M) or *Psm* ES4326/avrRpt2 (OD_600nm_ = 0.01). After 2 days, the systemic leaves were infiltrated with *Psm* ES4326 (OD_600nm_ = 0.001). Bacterial growth was measured 3 days after the second pathogen infection. Two leaf-discs (0.6 cm in diameter) were placed in one of the 8-strip tubes containing 1 metal bead and 500 μL sterilized 10 mM MgCl_2_ and grind with a SPEX^TM^ SamplePrep 2010 Geno/Grinder^TM^ (ThermoFisher Scientific) at 1500 strokes/min for 30 sec for twice. Then, the samples were centrifuged at 2000 rpm for 1 min. 10 × dilution (6 gradient dilutions per sample) was used for each sample. 10 μL aliquots from each dilution were spread on King’s B Medium. Allow the bacteria to grow for 2 days at 30 °C and then count individual colony number for each dilution. The bacteria number were calculated based on the dilution and the area of leaf discs.

For petiole exudate (PeX)-triggered plant defense, 3-week-old plants were first infiltrated with PeX. One days after, the treated leaves or distal leaves were infiltrated with *Psm* ES4326 (OD_600nm_ = 0.001) and bacterial growth was measured 3 days after pathogen infection.

### Luciferase imaging and circadian rhythm calculation

Independent T1 seedlings were grown on half strength of Murashige-Skoog (MS) media with 25 μg/mL hygromycin B (ThermoFisher Scientific) for 10 days at 22 °C under 12 hours light/12 hours dark condition. Then they were sprayed with 2.5 mM luciferin (Gold Biotechnology) in 0.02% Triton X-100 (Sigma) and transferred to constant light condition one day before imaging. Imagines were taken every 2 hours with exposure time of 20 min using a charge-coupled device camera (PIXIS 2048). The quantifications of bioluminescence intensity were performed using Image J. Periods of circadian rhythm were inferred from a sine wave as previously described (*64*).

### Chromatin immunoprecipitation (ChIP)

ChIP was performed as previously described (*65*) with minor modifications. Pierce^TM^ Anti-HA magnetic beads (ThermoFisher Scientific) were used for immunoprecipitation (IP) as described in the manufacturer’s protocol, and primes used are listed in Table S1.

### Protein S-sulfenylation and S-sulfinylation

Protein S-sulfenylation and S-sulfinylation were measured using a biotin-switch method as described previously (*66, 67*) with modifications. Leaf tissues (2 g) were collected from 3-week-old plants, ground in liquid nitrogen, and suspended in 25 mL HEN buffer [250 mM HEPES-NaOH (pH 7.7), 1 mM EDTA, 0.1 mM neocuproine] supplemented with 1 × protease inhibitor cocktail (Roche), 2 % SDS, and 50 mM MMTS. The sample was then incubated at 50 °C for 1 hour with vortex under dark condition and the supernatant was collected after centrifugation (4 °C, 7830 rpm for 1 hour). The sample was then precipitated and washed with acetone.

For detecting S-sulfenylation, the pellet was suspended with 2-5 mL HEN buffer with 1 % SDS and supplied with 2 mM biotin-HPDP and 50 mM sodium arsenite and incubated at 25 °C for 1 hour with vortex under dark condition.

For detecting S-sulfinylation, the pellet was suspended in 2-5 mL HEN buffer with 0.4 % SDS supplied with 20 mM DiaAlk (AOBIOUS) and incubated at 37 °C for 2 hours with vortex under dark condition. The solution is precipitated and washed with acetone and resuspended with HEN buffer supplied with 0.2 mM azide-biotin, 0.1 mM TBTA, and 1 mM CuSO_4_, and incubated at 25 °C for 2 hours with vortex under dark condition.

The subsequent steps were used for detecting both S-sulfenylation and S-sulfinylation. After setting aside 5-10 μL solutions as the input for western blot, the rest solution was precipitated and washed with acetone. The modified protein was pulled down with Pierce^TM^ streptavidin-agarose (ThermoFisher Scientific). The protein was dissolved with 50-100 μL 20 mM HEPES-NaOH buffer, supplied with 1 mM EDTA, 10 mM NaCl, and 100 mM 2-mercaptoethanol. The sample is used for western blot analysis after mixing with 2 × SDS-PAGE buffer and 1 mM DTT.

### Protein purification and electrophoretic mobility shift assay (EMSA)

The CHE protein fused with glutathione-S-transferase (GST) was expressed in *E. coli* strain BL21 and purified using Pierce^TM^ glutathione magnetic agarose (ThermoFisher Scientific). The purified protein was digested with PreScission Protease (APEXBIO) to remove GST tag and treated with 25 mM DTT or 50 μM H_2_O_2_ at 25 °C for 30 min, the treated protein was cleaned up using a Micro Bio-Spin^TM^ P-6 Gel Column (BIO-RAD), and then they were further subjected to 5 mM sodium (meta) arsenite (m-arsenite) treatment at 25 °C for 30 min (Water treatment was used as the control). The treated protein was cleaned up using a Micro Bio-Spin^TM^ P-6 Gel Column and used for EMSA. EMSA was performed using a LightShift^TM^ Chemiluminescent EMSA kit (ThermoFisher Scientific) following the manufacturer’s protocol, and DNA probes used were listed in Table S1.

### Hydrogen peroxide (H_2_O_2_) quantification

Three leaf discs (0.5 cm in diameter) were collected, grounded in liquid nitrogen, and suspended in 600 μl pre-cooled 20 mM phosphate buffer (pH 6.5) containing 0.02 % acetanilide. After centrifugation at 4 °C (16,200 g for 10 min), the supernatant was transferred to a new pre-cooled tube. To quantify concentration of H_2_O_2_, 50 μl of the solution was used according to the protocol provided by Amplex^TM^ Red Hydrogen Peroxide Assay kit (Invitrogen). For each measurement, a H_2_O_2_ standard curve (0 to 100 μM, each in a volume of 50 ul) were generated. Luminescence was captured using the Victor3 plate reader (PerkinElmer) with excitation at 531 nm and emission detection at 595 nm. The H_2_O_2_ concentration were determined using the standard curve and divided by the area of the leaf disc. For H_2_O_2_ collected from PeX, the concentration was determined directly according to the standard curve.

### Live imaging of reactive oxygen species (ROS)

Live ROS imaging was recorded as previously described (*51*) with modifications. 3-week-old plant leaves were infiltrated with 50 μM 2’,7’-dichlorodihydrofluorescein diacetate (H_2_DCFDA) (Millipore-Sigma) in 50 mM phosphate buffer (pH 7.4) and allowed for 6-8 hours to recover. Then the infiltrated plants were treated with pathogen or mock solution, and images were taken every 20 min for 2 days using microscope (LEICA M205 FA). Imaging data were analyzed with ImagJ (FIJI).

### Statistical analysis

Two-tailed Student’s t-tests and two-way ANOVA were performed using Graphpad prism 8. Sample size is described in the relevant figure legends and experiments have been performed at least twice with similar results. For SA and Pip measurement, CHIP-qPCR and luciferase imaging, at least 3 independent samples were used in the figures.

---

**Fig. S1.**
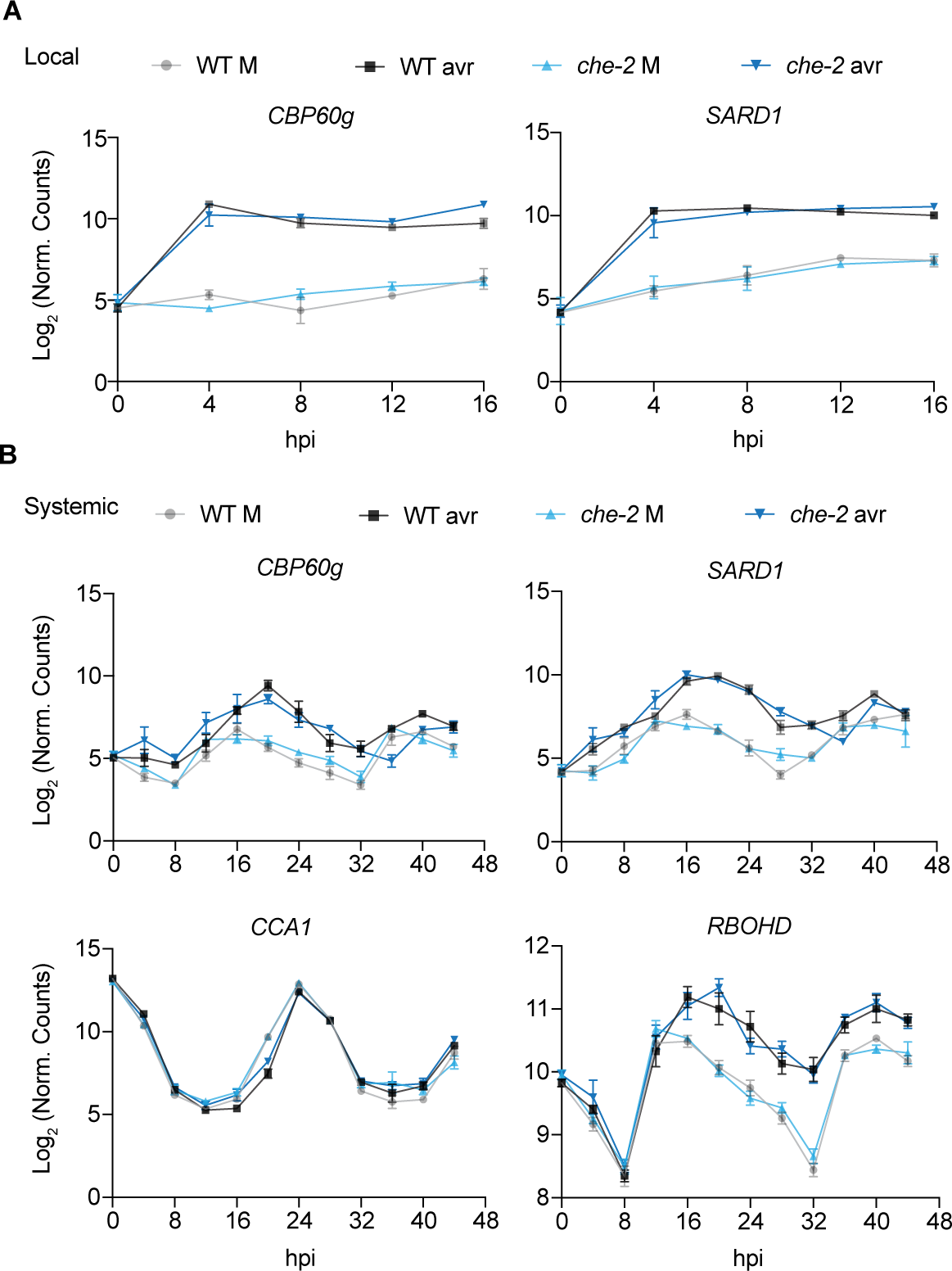
Normalized read counts from RASL-seq. (**A** and **B**) Normalized transcript read counts in WT and *che-2* plants collected from local (A) and systemic (B) tissues after mock (M; 10 mM MgCl_2_) or *Psm* ES4326/avrRpt2 (avr; OD_600nm_ = 0.01) treatment. Data are means ± SEMs (*n* ≥ 3). hpi, hours post infiltration.

**Fig. S2.**
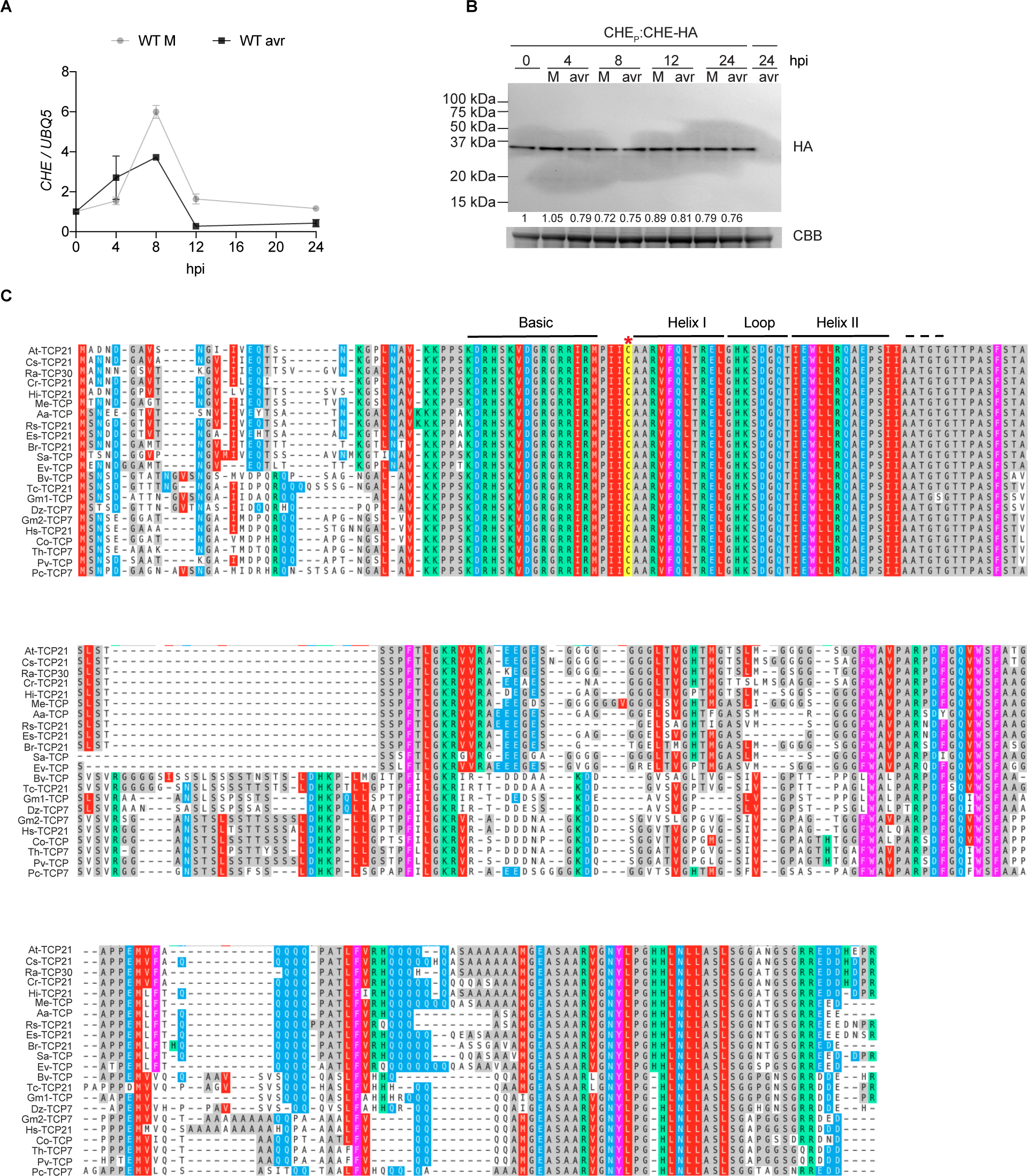
CHE transcript and protein levels and conservation of the cysteine residue in CHE among plant species. (**A** and **B**) CHE transcript (A) and protein level (B) changes in systemic tissues after local mock (M; 10 mM MgCl_2_) or *Psm* ES4326/avrRpt2 (avr; OD_600nm_ = 0.01) treatment. hpi, hours post infiltration. Data are means ± SEMs. *n* = 3 for (A). (**C**) Sequence alignment of CHE homologs in plants. The red star indicates the conserved cysteine residue. *Arabidopsis thaliana* (*At*), *Camelina sativa* (*Cs*), *Rorippa aquatica* (*Ra*), *Capsella rubella* (*Cr*), *Hirschfeldia incana* (*Hi*), *Microthlaspi erraticum* (*Me*), *Arabis alpina* (*Aa*), *Raphanus sativus* (*Rs*), *Eutrema salsugineum* (*Es*), *Brassica rapa* (*Br*), *Sinapis alba* (*Sa*), *Eruca vesicaria* (*Ev*), *Bauhinia variegate* (*Bv*), *Theobroma cacao* (*Tc*), *Gossypium mustelinum* (*Gm*), *Durio zibethinus* (*Dz*), *Glycine max* (*Gm*), *Hibiscus syriacus* (*Hs*), *Corchorus olitorius* (*Co*), *Tarenaya hassleriana* (*Th*), *Phaseolus vulgaris* (*Pv*), *Prosopis cineraria* (*Pc*).

**Fig. S3.**
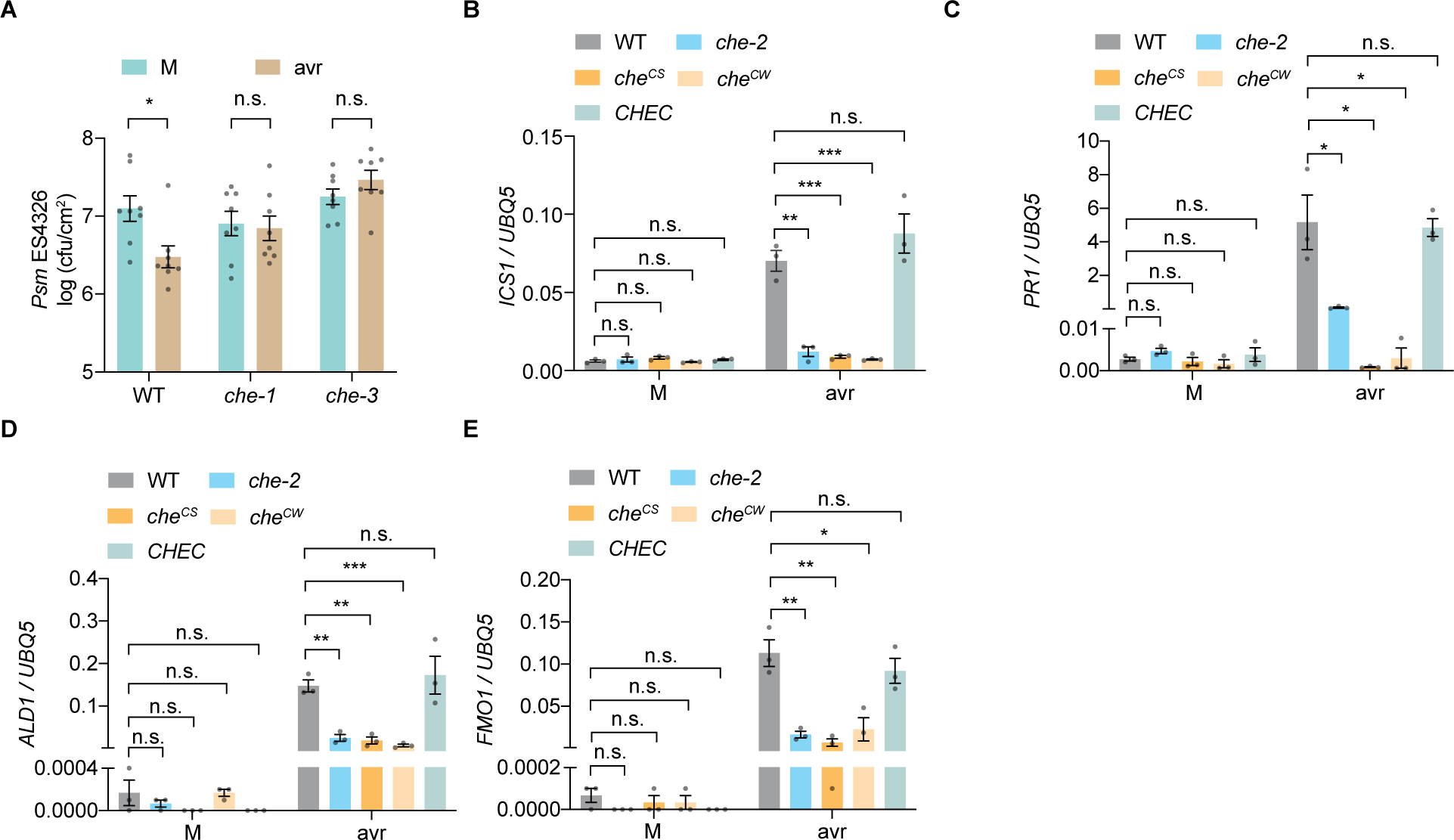
Bacterial growth and defense-related gene expression in systemic tissues. (**A**) Bacterial growth after pathogen challenge. Plants were infiltrated with mock (M; 10 mM MgCl_2_) or *Psm* ES4326/avrRpt2 (avr; OD_600nm_ = 0.01) 2 days before infiltration of the systemic tissues with *Psm* ES4326 (OD_600nm_ = 0.001), and bacterial growth was measured 3 days after the second infiltration. Data are means ± SEMs (*n* = 8). (**B**-**E**) Transcriptional levels of *ICS1* (B), *ALD1* (C), *FMO1* (D) and *PR1* (E) in systemic tissues after local M or avr treatment. Data are means ± SEMs (*n* = 3). *CHEC*, *che-2* transformed with the WT *CHE* expressed by its native promoter. *che^CS^* and *che^CW^*, transformants expressing the cysteine-to-serine and cysteine-to-tryptophan mutant *che*, respectively. Significant differences were calculated using two-tailed Student’s t-tests. \*\*\**P* < 0.001; \*\**P* < 0.01; \**P* < 0.05; n.s., not significant.

**Fig. S4.**
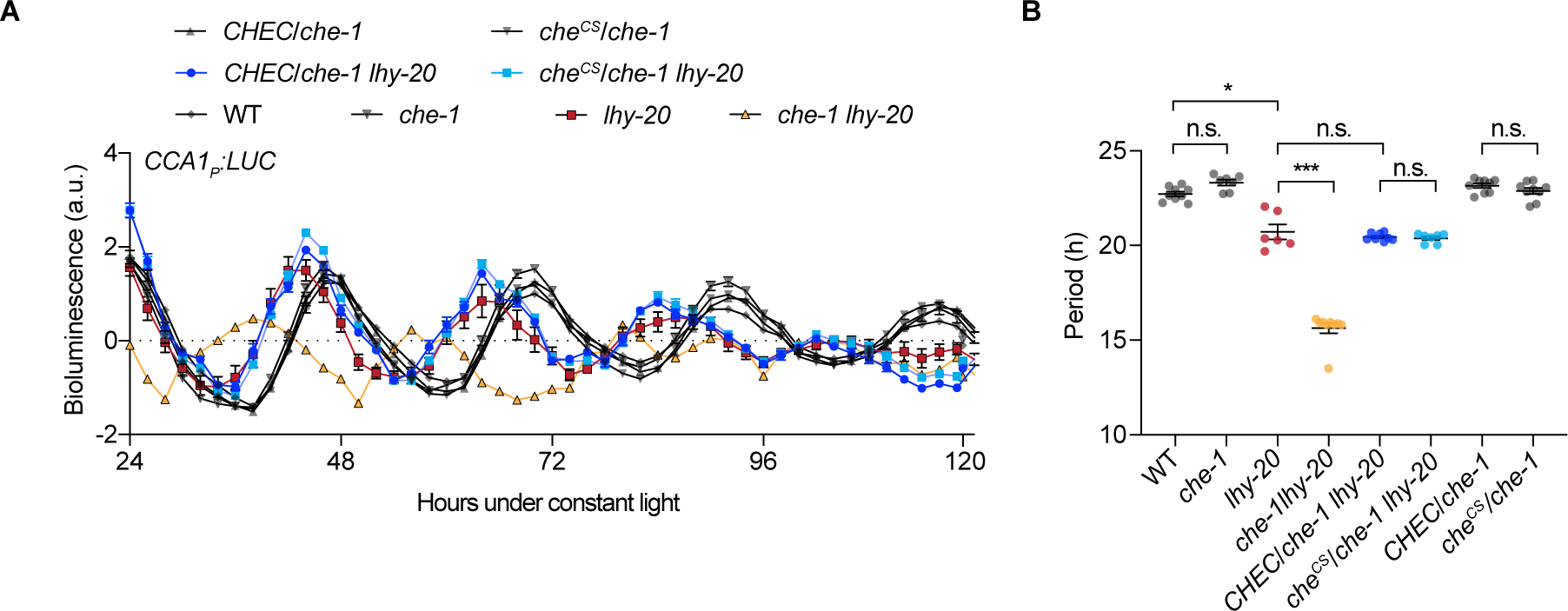
The cysteine mutant can rescue the phenotype of *che-1* in the regulation of *CCA1*. (**A**) Bioluminescence activity of *CCA1_P_:LUC* under constant light condition. Data are means ± SEMs (*n* ≥ 6). *CHEC/che-1*, *che-1* transformed with the WT *CHE* expressed by its native promoter. *che^CS^*, transformants expressing the cysteine-to-serine mutant of CHE. (**B**) Period estimates of *CCA1_P_:LUC* activity. Data are means ± SEMs (*n* ≥ 6). Significant differences were calculated using two-tailed Student’s t-tests. \*\*\**P* < 0.001; \*\**P* < 0.01; \**P* < 0.05; n.s., not significant.

**Fig. S5.**
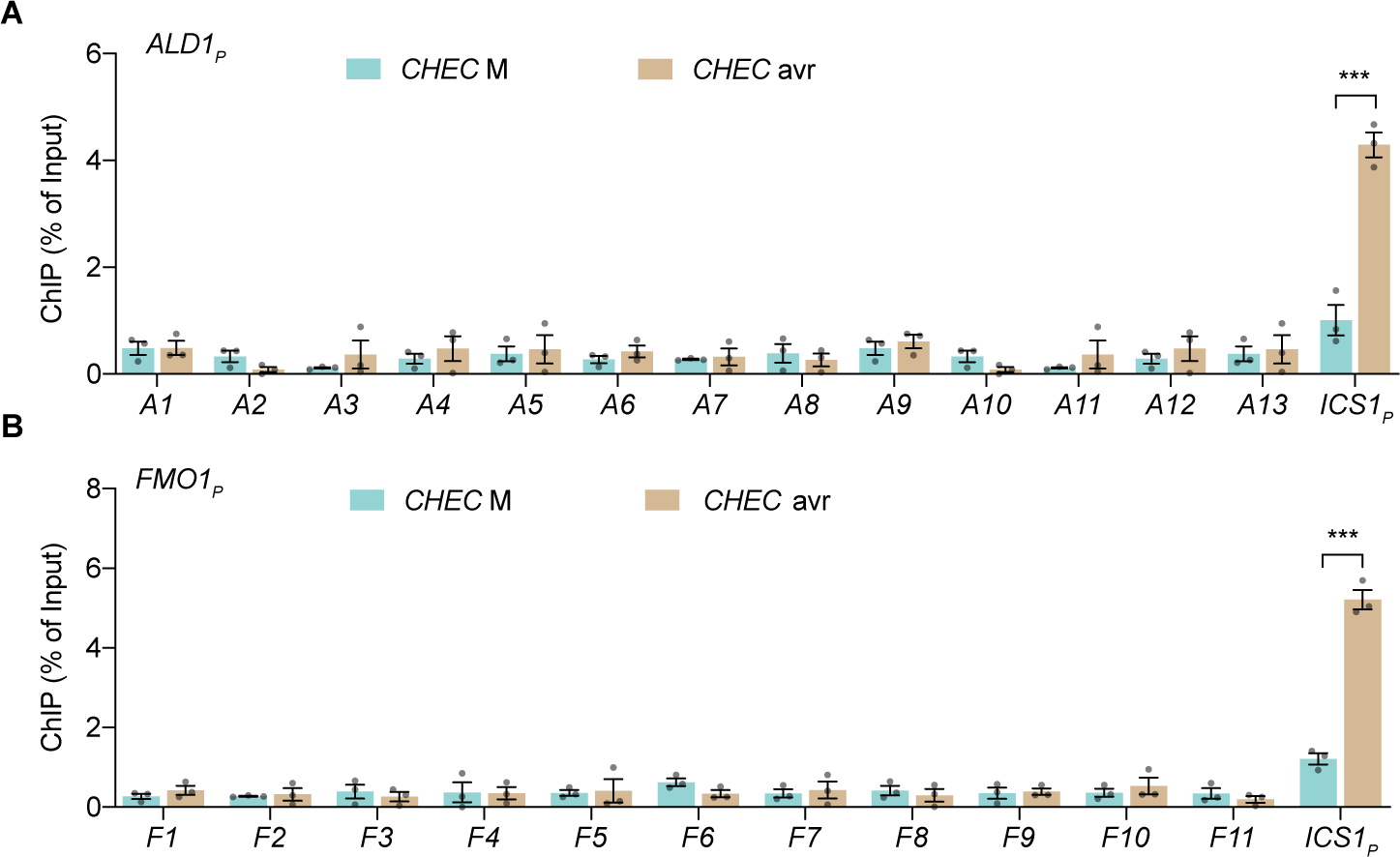
CHE does not bind to the *ALD1* or *FMO1* promoter. (**A** and **B**) ChIP-qPCR analysis of CHE binding to the *ALD1* promoter (*ALD1_P_*) (A) and the *FMO1* promoter (*FMO1_P_*) (B) in systemic tissues 24 hours after local mock (M; 10 mM MgCl_2_) or *Psm* ES4326/avrRpt2 (avr; OD_600nm_ = 0.01) treatment. *A1* to *A13*, qPCR fragments covering the promoter sequence of *ALD1* (1908 bp upstream ATG to 46 bp downstream ATG); *F1* to *F11*, qPCR fragments covering the promoter sequence of *FMO1* (1583 bp upstream ATG to 53 bp downstream ATG); *ICS1_P_*, the *ICS1* promoter sequence carrying the TCP-binding site. Data are means ± SEMs (*n* = 3). Significant differences were calculated using two-tailed Student’s t-tests. \*\*\**P* < 0.001.

**Fig. S6.**
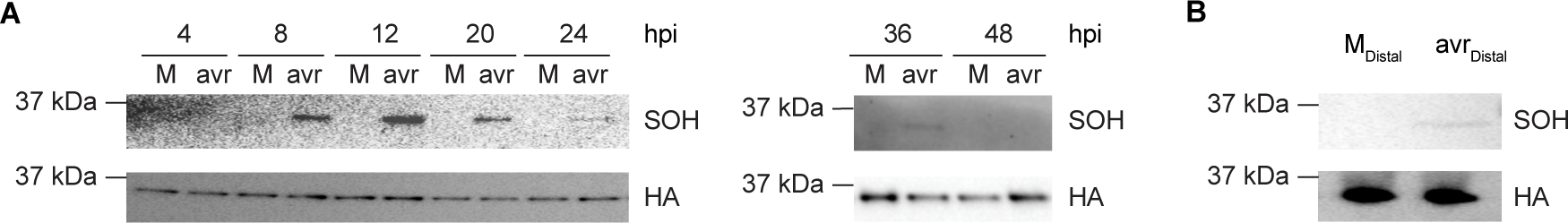
*In vivo* sulfenylation of CHE. (**A**) Time-course sulfenylation (SOH) of CHE in systemic tissues after local mock (M; 10 mM MgCl_2_) or *Psm* ES4326/avrRpt2 (avr; OD_600nm_ = 0.01) treatment. hpi, hours post infiltration. (**B**) Sulfenylation of CHE in the distal tissues 24 hours after treatment of the lower leaves with M or avr.

**Fig. S7.**
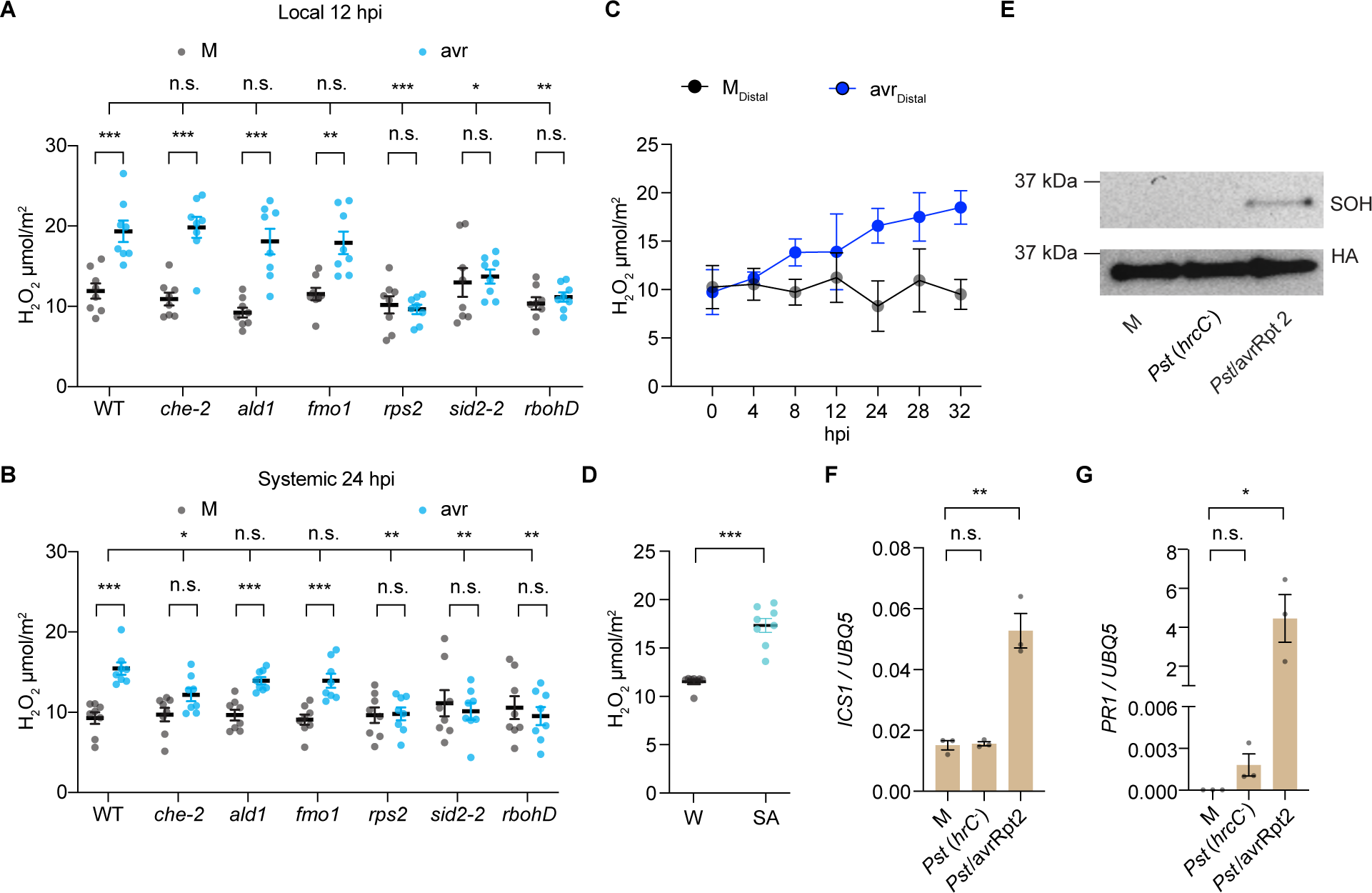
H_2_O_2_ is the initial mobile signal for ETI-induced SAR. (**A** and **B**) Levels of H_2_O_2_ produced in the local tissues 12 hours after mock (M; 10 mM MgCl_2_) or *Psm* ES4326/avrRpt2 (avr; OD_600nm_ = 0.01) treatment (A) or in the systemic tissues 24 hours after the local treatment (B). hpi, hours post infiltration. Data are means ± SEMs (*n* = 8). (**C**) Time-course H_2_O_2_ production in the distal tissues when lower leaves were treated with M or avr. Data are means ± SEMs (*n* = 5). (**D**) Levels of H_2_O_2_ produced 24 hours after water (W) or 1 mM SA treatment in WT plants. Data are the means ± SEMs (*n* = 8). (**E**-**G**) Sulfenylation of CHE (E) and expressions of *ICS1* (F) and *PR1* (G) in systemic tissues after local treatment with M, *Pst* DC3000 *hrcC^-^* [*Pst* (*hrcC^-^*)] or *Pst* DC3000/avrRpt2 (*Pst*/avrRpt2). Data are means ± SEMs (*n* = 3). Significant differences were calculated using either two-tailed Student’s t-tests or two-way ANOVA. \*\*\**P* < 0.001; \*\**P* < 0.01; \**P* < 0.05; n.s., not significant.

**Fig. S8.**
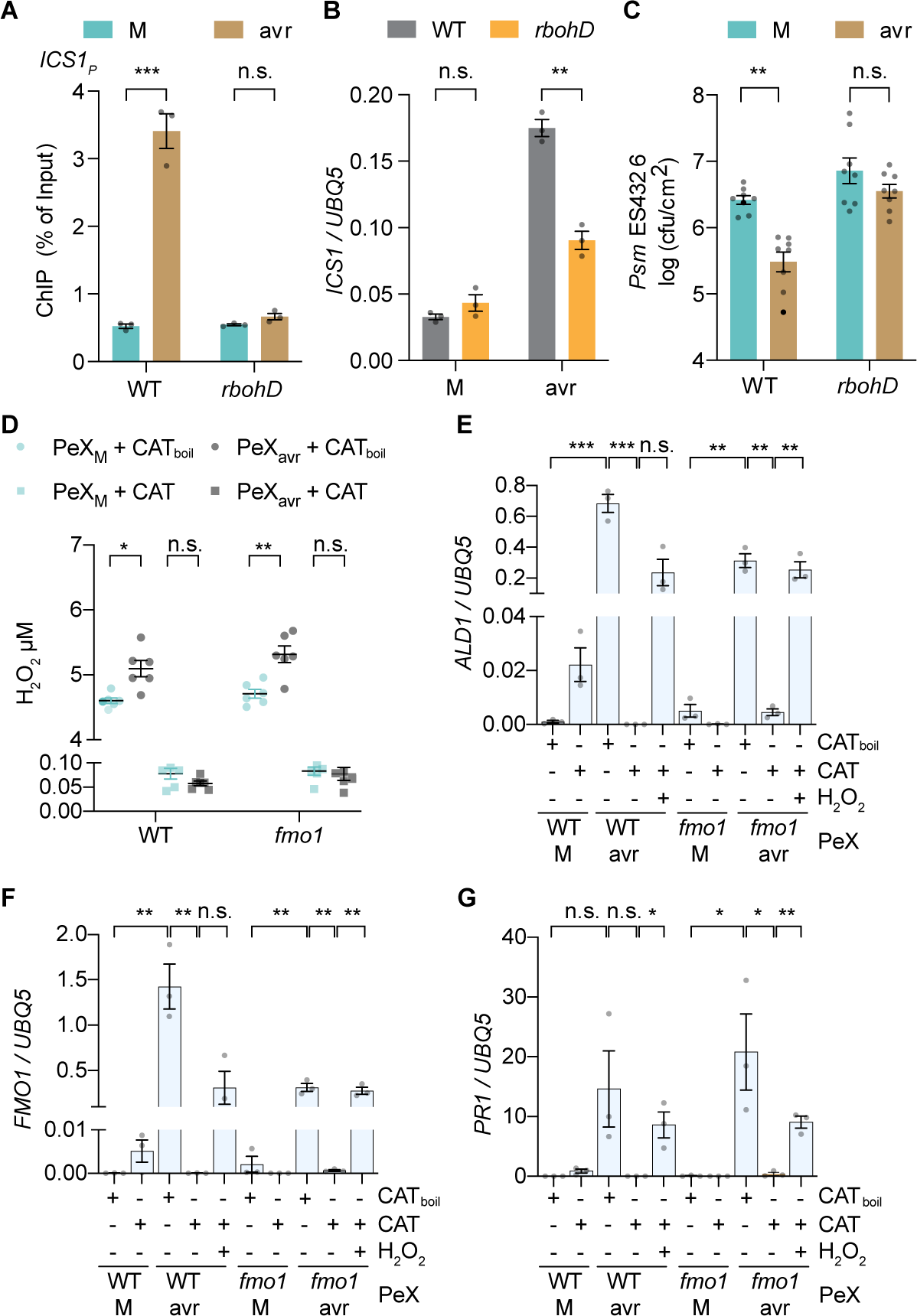
H_2_O_2_ produced by RBOHD is the signal that induces systemic SA synthesis and Pip accumulation to confer SAR. (**A**-**C**) ChIP-qPCR analysis of CHE binding to the *ICS1* promoter carrying the TCP-binding site (*ICS1_P_*) (A), transcriptional level of *ICS1* (B) and bacterial growth (C) in systemic tissues of WT or *rbohD* plants after local mock (M; 10 mM MgCl_2_) or *Psm* ES4326/avrRpt2 (avr; OD_600nm_ = 0.01) treatment. Data are means ± SEMs. *n* = 3 for (A and B), *n* = 8 for (C). *ICS1_P_*, the *ICS1* promoter sequence carrying the TCP-binding site. (**D**) Levels of H_2_O_2_ in petiole exudates (PeX) collected from WT and *fmo1* plants after treatment with catalase (CAT) or heat-denatured catalase (CAT_boil_). Data are means ± SEMs (*n* = 6). (**E**-**G**) The expression of *ALD1* (E), *FMO1* (F) and *PR1* (G) after infiltration with PeX collected from WT or *fmo1* treated with M or avr. Data are means ± SEMs (*n* = 3). Significant differences were calculated using two-tailed Student’s t-tests. \*\*\**P* < 0.001; \*\**P* < 0.01; \**P* < 0.05; n.s., not significant.

## Notes

### Summary of Updates

Updated conflict of interest and copyright.

